# DeepSpot2Cell: Predicting Virtual Single-Cell Spatial Transcriptomics from H&E images using Spot-Level Supervision

**DOI:** 10.1101/2025.09.23.678121

**Authors:** Kalin Nonchev, Glib Manaiev, Viktor H Koelzer, Gunnar Rätsch

## Abstract

Spot-based spatial transcriptomics (ST) technologies like 10x Visium quantify genome-wide gene expression and preserve spatial tissue organization. However, their coarse spot-level resolution aggregates signals from multiple cells, preventing accurate single-cell analysis and detailed cellular characterization. Here, we present DeepSpot2Cell, a novel DeepSet neural network that leverages pretrained pathology foundation models and spatial multi-level context to effectively predict virtual single-cell gene expression from histopathological images using spot-level supervision. DeepSpot2Cell substantially improves gene expression correlations on a newly curated benchmark we specifically designed for single-cell ST deconvolution and prediction from H&E images. The benchmark includes 20 lung, 7 breast, and 2 pancreatic cancer samples, across which DeepSpot2Cell outperformed previous super-resolution methods, achieving respective improvements of 46%, 65%, and 38% in cell expression correlation for the top 100 genes. We hope that DeepSpot2Cell and this benchmark will stimulate further advancements in virtual single-cell ST, enabling more precise delineation of cell-type-specific expression patterns and facilitating enhanced downstream analyses.

**Code availability:** https://github.com/ratschlab/DeepSpot2Cell

## 1 Introduction

Spatial transcriptomics (ST) provides valuable insights into the spatial heterogeneity of tissue microenvironments and the mechanisms underlying disease progression [1, 2]. Despite its transformative potential, current ST technologies involve an inherent trade-off between spatial resolution and transcriptome coverage [3]. For example, 10x Visium captures the whole transcriptome but at a coarse spot-level resolution, integrating signals from 1–10 cells depending on cell size [4]. As a result, each spot represents a bulk of cells, making it difficult to distinguish individual cell types or states. In contrast, 10x Xenium achieves single-cell resolution but is restricted to a targeted gene panel, which may exclude relevant biomarkers. Newer technologies have advanced spatially resolved single-cell expression profiling by enabling subcellular, full-transcriptome coverage (e.g., 10x Visium HD). However, these methods still face major challenges [5], including low gene-detection sensitivity, high error rates, and high costs, which limit their applicability in clinical studies. Achieving true single-cell resolution across all protein-coding genes would allow for more precise cellular annotations and a detailed, multi-modal view of biological mechanisms and cellular interactions.

At the same time, data generated using the more established spot-level Visium technology are rapidly accumulating in public databases [6, 7, 8] and are supporting the first large-scale cohorts (e.g., 7,000 patients in MOSAIC [9]). Consequently, robust deconvolution algorithms are urgently needed to accurately reconstruct single-cell transcriptomic profiles for high-resolution cellular analysis.

Recently, advances in deep learning have demonstrated that hematoxylin and eosin (H&E)-stained histological images can be used to effectively predict ST profiles [10, 11, 12, 13, 14] with some genes exceeding a correlation of 0.60 [14]. These methods represent a promising, cost-effective, and scalable alternative to conventional sequencing techniques. Building on this, early studies have explored the prediction of super-resolution transcriptomic data [15, 16], which produce superpixel-level expression maps rather than precise cell-level profiles. Despite these advances, achieving true single-cell transcriptomic resolution remains a major challenge.

To this end, we present DeepSpot2Cell, a novel deep learning model that leverages recent pathology foundation models alongside spatial multi-level context to accurately predict virtual single-cell gene expression from H&E images using spot-level supervision. DeepSpot2Cell uses a permutation-invariant DeepSet architecture to represent spots as bags of individual cells, enabling the model to learn each cell’s contribution during training and perform single-cell prediction at inference.

We evaluate our model on a newly curated benchmark, which we specifically designed to assess two fundamental tasks: deconvolving retrospective ST datasets and predicting single-cell expression using unseen H&E images. The benchmark includes 20 lung, 7 breast, and 2 pancreatic cancer samples, sequenced at single-cell resolution with 10x Xenium and structured to mimic 10x Visium spot-level data. The results demonstrate a substantially improved reconstruction of single-cell transcriptomic profiles compared to previous super-resolution models, with consistent performance validated across in-sample, out-of-sample, and out-of-distribution scenarios.

To our knowledge, DeepSpot2Cell is the first deep learning model to leverage pathology foundation models to predict virtual single-cell ST from H&E images using spot-level supervision. This strategy takes advantage of the rapidly growing spot-level ST datasets by learning robust single-cell mappings between tissue images and transcriptomic profiles. It enables both the augmentation of retrospective ST cohorts with single-cell resolution and the annotation of histopathological images with virtual single-cell ST, thereby supporting more precise and detailed cellular analyses for biomedical research.

## 2 Related Works

### 2.1 Pathology foundation models

Pathology foundation models (PFM) are trained on large-scale histopathology datasets using self-supervised techniques such as contrastive learning or masked image modeling. Notable examples include UNI [17], Phikon-v2 [18], and H-Optimus-0 [19], which mostly rely on vision transformers (ViT) to learn high-dimensional morphological representations. These models achieve state-of-the-art performance across a range of computational pathology tasks [20, 21].

### 2.2 Spatial transcriptomics prediction from H&E images

ST sequencing methods generate spatially resolved transcriptomic profiles aligned with H&E images. For example, on the 10x Genomics Visium platform, each spot covers 55µm of tissue area and captures transcripts from 1-10 cells depending on the cell size [4].

With the increasing availability of such molecular datasets [6, 7, 8], specialized machine-learning models have been developed to predict ST from H&E images using convolutional neural networks [10], vision transformers [12, 13], or contrastive learning [11]. The recent DeepSpot model [14] introduces two key innovations: leveraging a PFM to extract informative spot representations and integrating spatial multi-level tissue and neighborhood context. This enables the prediction of 5,000 genes with substantially higher accuracy and up to six-fold greater coverage than previous models.

### 2.3 Super-resolution-based deconvolution models from H&E images

Super-resolution methods aim to improve the spatial resolution of ST by integrating H&E images. For example, iStar [22] uses hierarchical vision transformers to extract histology features at a 16×16-pixel scale, capturing fine-grained tissue characteristics. scstGCN [23] combines graph convolutional networks with PFM and spatial information to capture the relationships among adjacent superpixels. However, these methods output high-dimensional superpixel expression maps rather than precise cell transcriptomic profiles, requiring custom post-processing to approximate individual cell expression.

## 3 Methods

### 3.1 Model architecture

DeepSpot2Cell extends the DeepSets architecture [24] to integrate spatial and multi-tissue context from histopathology images for accurate cell expression prediction using spot-level supervision. As illustrated in Figure 1, for each cell *j* within spot *i*, the model extracts features from three H&E image inputs using frozen PFM: (i) the cropped cell tile defined by the segmentation mask 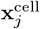, (ii) the full spot tile containing the cell 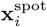, and (iii) the neighboring spot tile(s)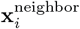

**Figure 1:**
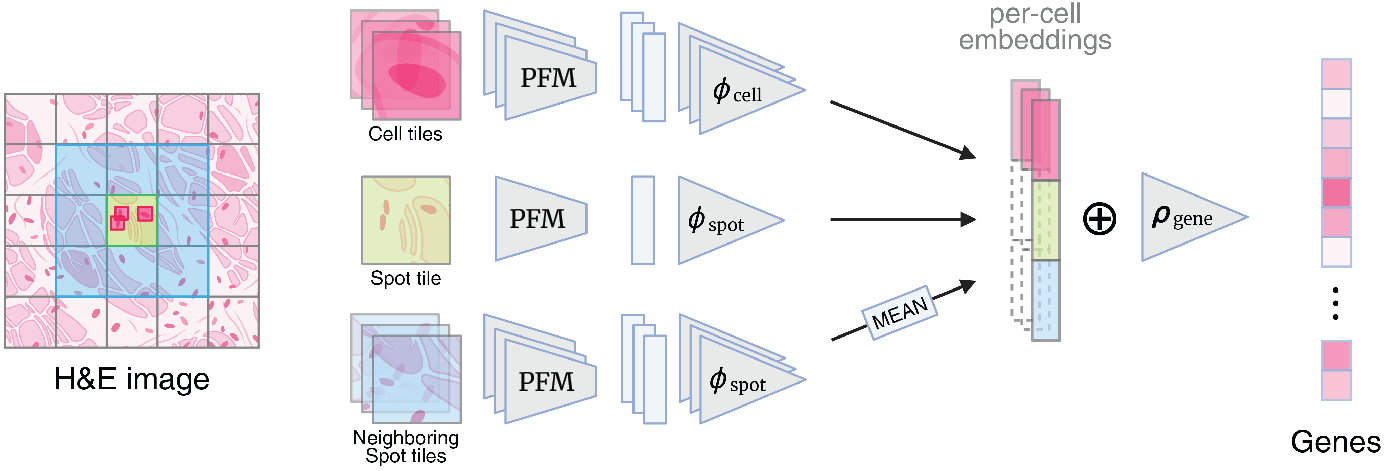
DeepSpot2Cell predicts virtual single-cell spatial transcriptomics as follows: (1) During training, the model takes as input (i) the cropped cell tile defined by the segmentation mask, (ii) the full spot tile containing the cell, and (iii) the neighboring spot tile(s). All tiles are first processed through a pathology foundation model (PFM) before being used to train the model to regress spot-level gene expression; (2) During inference, the model takes as input only the cell tile of interest along with (ii) and (iii), again after PFM processing, and predicts the virtual transcriptomic profile at the cell level.

**Figure 2:**
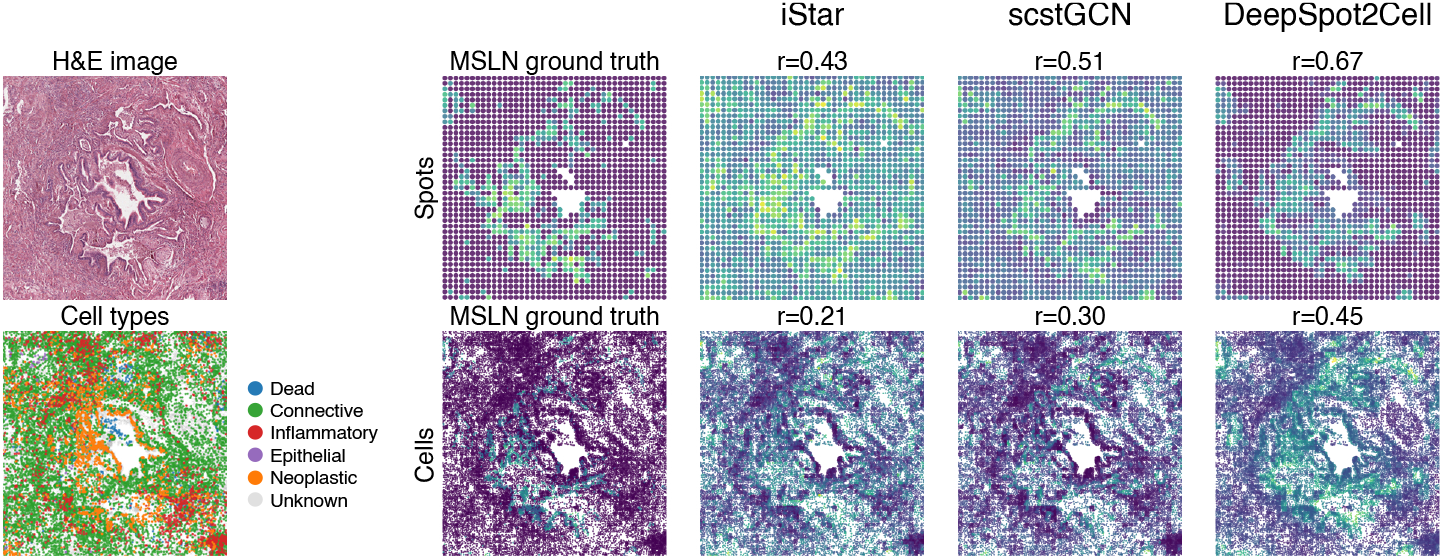
H&E image and CellViT [26] cell-type annotations for slide NCBI867 (lung dataset). Annotations and model predictions of *MSLN* expression across spots and cells.

**Figure 3:**
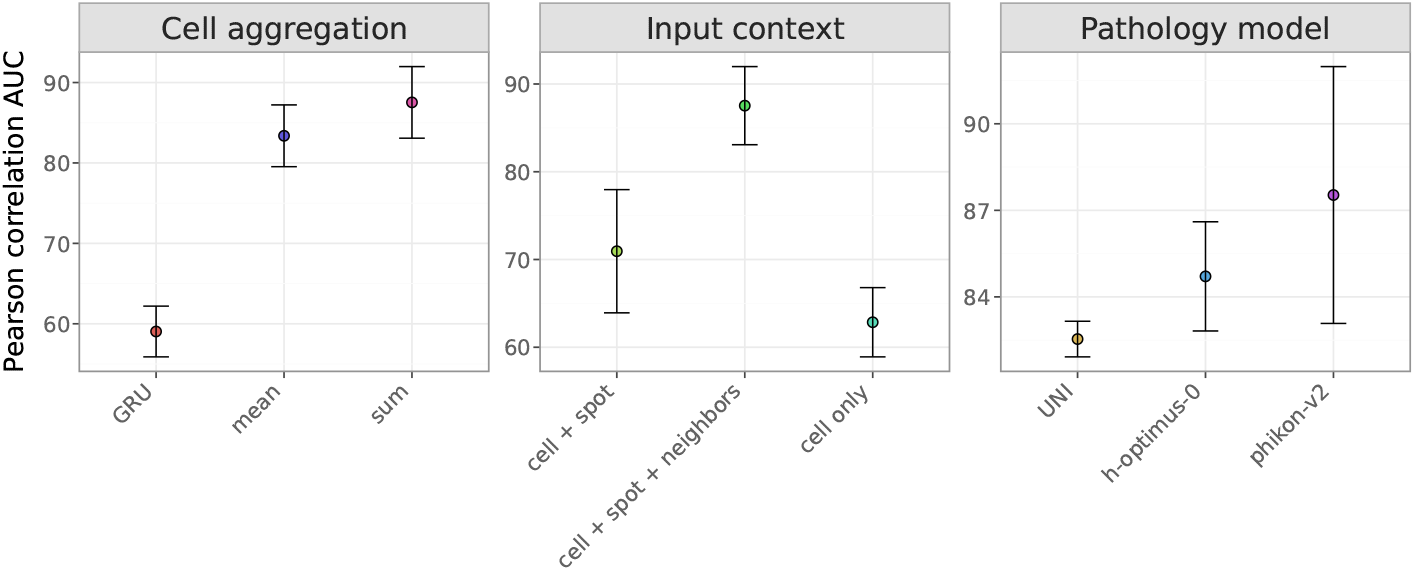
Different DeepSpot2Cell components compared based on the area under the Pearson correlation gene curve computed on the cells from within spots (IS_*in*_).

For each cell, PFM embeddings are processed via dedicated two-layer multilayer perceptrons (MLP): *ϕ*_cell_ for cell tiles, and *ϕ*_spot_ for spot and neighboring contexts. These embeddings are concatenated to form the integrated cell representation:

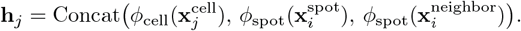

Cell embeddings within each spot are aggregated by summation, ensuring permutation invariance and accommodating variable cell counts within a spot:

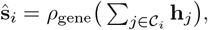

where *ρ*_gene_ is a two-layer MLP gene prediction head generating the predicted gene expression vector ŝ_*i*_ ∈ ℝ^*G*^ for spot *i*, and 𝒞_*i*_ denotes the set of cells within that spot. Notably, the summation aggregation naturally models transcript count additivity and ensures robustness to cell order permutations.

### 3.2 Benchmarking of virtual cell transcriptomic profiles inferred from H&E images

10x Visium measures gene expression at the spot level, but single-cell resolution is needed for accurate evaluation of prediction methods. To address this, we gathered 10x Xenium datasets across multiple cancer and tissue types (Table 2, Figure 4) with true single-cell resolution to establish a newly dedicated benchmark. Building upon the HEST-1k benchmark [20], we derived pseudo spot-level gene expression profiles by aggregating single-cell transcript counts within each 55µm spatial spot, consistent with 10x Visium. To account for cell size variability [25], a cell was considered fully contained within a spot if its nucleus was located at least 10µm inside the spot boundary.

**Figure 4:**
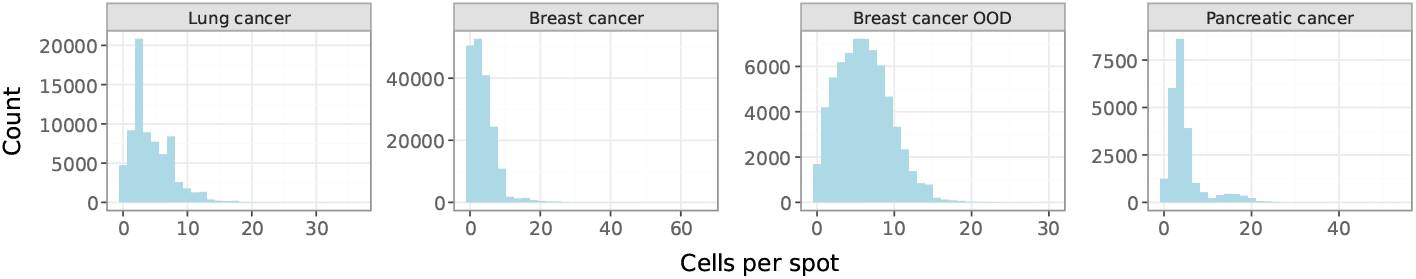
Distribution of cell counts per spot across datasets.

#### Evaluation

Models are trained exclusively on spot-level data, and their performance is assessed by comparing predicted gene expression to single-cell ground truth using per-gene Pearson correlation. Two key evaluation tasks were considered: deconvolution of in-sample (IS) cells in spots seen during training, and prediction of single-cell expression for out-of-sample (OOS) cells from unseen samples within the cohort, and out-of-distribution (OOD) cells from samples belonging to a different cohort.

## 4 Experiments

### 4.1 DeepSpot2Cell enables cell expression deconvolution from spatial transcriptomics spots

Table 1 summarizes the performance of a two-layer MLP baseline, previous super-resolution-based models (iStart and scstGCN), and DeepSpot2Cell in deconvolving spot-based ST from H&E images. The in-sample (IS) scenario benchmarks deconvolution performance. IS is further subdivided into IS_*in*_, assessing gene correlation among cells within spots, and IS_*out*_, assessing cells outside spots.

**Table 1:** Benchmark of single-cell expression prediction across lung, breast, and pancreatic cancer datasets. Average Pearson correlation between predicted and ground-truth single-cell gene expression is reported for the top 100 most predictive genes.

| Model | Lung cancer (n=20) |  |  | Breast cancer (n=7) |  |  |  | Pancreatic cancer (n=2) |  |  |
| --- | --- | --- | --- | --- | --- | --- | --- | --- | --- | --- |
| | $IS_{in}$ | $IS_{out}$ | OOS | $IS_{in}$ | $IS_{out}$ | OOS | OOD | $IS_{in}$ | $IS_{out}$ | OOS |
| MLP | 0.20<br>(0.01) | 0.24<br>(0.00) | 0.19<br>(0.00) | 0.30<br>(0.01) | 0.34<br>(0.01) | 0.14<br>(0.00) | 0.25<br>(0.01) | 0.18<br>(0.00) | 0.09<br>(0.00) | 0.10<br>(0.00) |
| iStar | 0.28<br>(0.01) | 0.28<br>(0.00) | 0.15<br>(0.00) | 0.34<br>(0.01) | 0.25<br>(0.01) | 0.10<br>(0.00) | 0.17<br>(0.00) | 0.28<br>(0.01) | 0.13<br>(0.00) | 0.10<br>(0.00) |
| scstGCN | 0.32<br>(0.01) | 0.35<br>(0.01) | 0.24<br>(0.01) | 0.37<br>(0.01) | 0.33<br>(0.01) | 0.17<br>(0.00) | 0.24<br>(0.01) | 0.29<br>(0.01) | 0.14<br>(0.00) | 0.08<br>(0.00) |
| DeepSpot2Cell (ours) | **0.39**<br>(0.01) | **0.41**<br>(0.01) | **0.35**<br>(0.01) | **0.43**<br>(0.01) | **0.41**<br>(0.01) | **0.28**<br>(0.01) | **0.37**<br>(0.01) | **0.32**<br>(0.01) | **0.16**<br>(0.00) | **0.11**<br>(0.00) |

$IS_{in}$ : gene correlation among cells within spots. $IS_{out}$ : gene correlation among cells outside spots. OOS: gene correlation among cells on hold-out patients, same cohort. OOD: gene correlations on cells from different cohort.

**Table 2:** HEST-1k datasets used in this study.

| Organ | Split | Dataset | Samples | Spots | Cells |
| --- | --- | --- | --- | --- | --- |
| Lung | Training | Image-based spatial transcriptomics identifies molecular niche dysregulation associated with distal lung remodeling in pulmonary fibrosis [33] | 20 | 76,023 | 998,856 |
| Breast | Training | High-resolution mapping of the tumor microenvironment using integrated single-cell, spatial and in-situ analysis [28] | 3 | 26,568 | 415,320 |
|  | Training | FFPE human breast using the entire sample area [29] | 2 | 177,885 | 1,766,768 |
|  | OOD Eval. | FFPE human breast with pre-designed panel [30] | 2 | 65,944 | 1,144,523 |
| Pancreas | Training | Pancreatic cancer with Xenium human multi-tissue and cancer panel [31] | 1 | 20,842 | 189,736 |
|  | Training | FFPE human pancreas with Xenium multimodal cell segmentation [32] | 1 | 5,827 | 139,581 |

DeepSpot2Cell substantially improved the single-cell gene expression deconvolution across the three cancer datasets. For example, in the lung cancer dataset, DeepSpot2Cell increased the IS_*in*_ Pearson correlation across the top 100 genes by 22%, from 0.32 (best competitor, scstGCN) to 0.39, with some genes exceeding a correlation of 0.50 (Figure 5). Notably, DeepSpot2Cell’s transcriptomic predictions for cells located within spots (IS_*in*_) and outside of spots (IS_*out*_) are similarly accurate in lung and breast cancer, indicating that the model does not overfit to the spot-level signals (Table 3, 4).

**Table 3:**
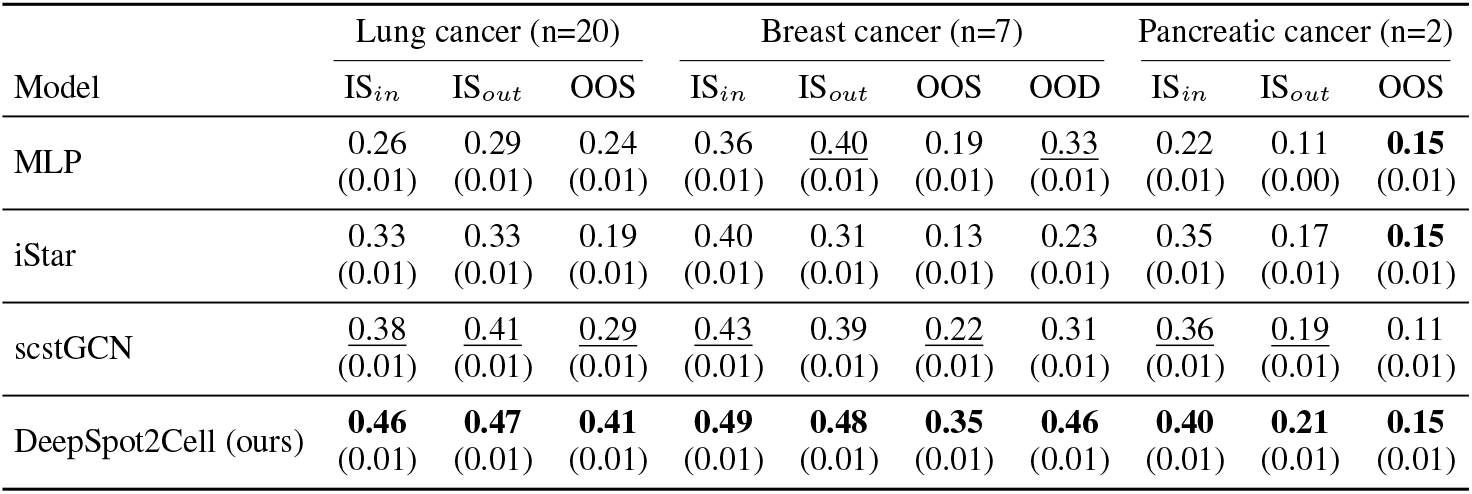
Benchmark of single-cell expression prediction across lung, breast, and pancreatic cancer datasets. Average Pearson correlation between predicted and ground-truth single-cell gene expression is reported for the top 50 most predictive genes.

**Table 4:** Benchmark of single-cell expression prediction across lung, breast, and pancreatic cancer datasets. Average Pearson correlation between predicted and ground-truth single-cell gene expression is reported for the top 200 most predictive genes.

| Model | Lung cancer (n=20) |  |  | Breast cancer (n=7) |  |  |  | Pancreatic cancer (n=2) |  |  |
| --- | --- | --- | --- | --- | --- | --- | --- | --- | --- | --- |
| | $IS_{in}$ | $IS_{out}$ | OOS | $IS_{in}$ | $IS_{out}$ | OOS | OOD | $IS_{in}$ | $IS_{out}$ | OOS |
| MLP | 0.13<br>(0.00) | 0.17<br>(0.00) | 0.14<br>(0.00) | 0.23<br>(0.00) | <u>0.25</u><br>(0.00) | 0.09<br>(0.00) | <u>0.17</u><br>(0.00) | 0.13<br>(0.00) | 0.06<br>(0.00) | **0.07**<br>(0.00) |
| iStar | 0.21<br>(0.00) | 0.22<br>(0.00) | 0.11<br>(0.00) | 0.26<br>(0.00) | 0.17<br>(0.00) | 0.05<br>(0.00) | 0.11<br>(0.00) | <u>0.21</u><br>(0.00) | 0.08<br>(0.00) | <u>0.06</u><br>(0.00) |
| scstGCN | <u>0.24</u><br>(0.00) | <u>0.28</u><br>(0.00) | <u>0.18</u><br>(0.00) | <u>0.29</u><br>(0.00) | 0.24<br>(0.00) | <u>0.12</u><br>(0.00) | 0.16<br>(0.01) | 0.20<br>(0.00) | <u>0.09</u><br>(0.00) | 0.04<br>(0.00) |
| DeepSpot2Cell (ours) | **0.32**<br>(0.00) | **0.33**<br>(0.00) | **0.29**<br>(0.00) | **0.35**<br>(0.00) | **0.32**<br>(0.00) | **0.19**<br>(0.00) | **0.26**<br>(0.01) | **0.24**<br>(0.00) | **0.10**<br>(0.00) | <u>0.06</u><br>(0.00) |

**Figure 5:**
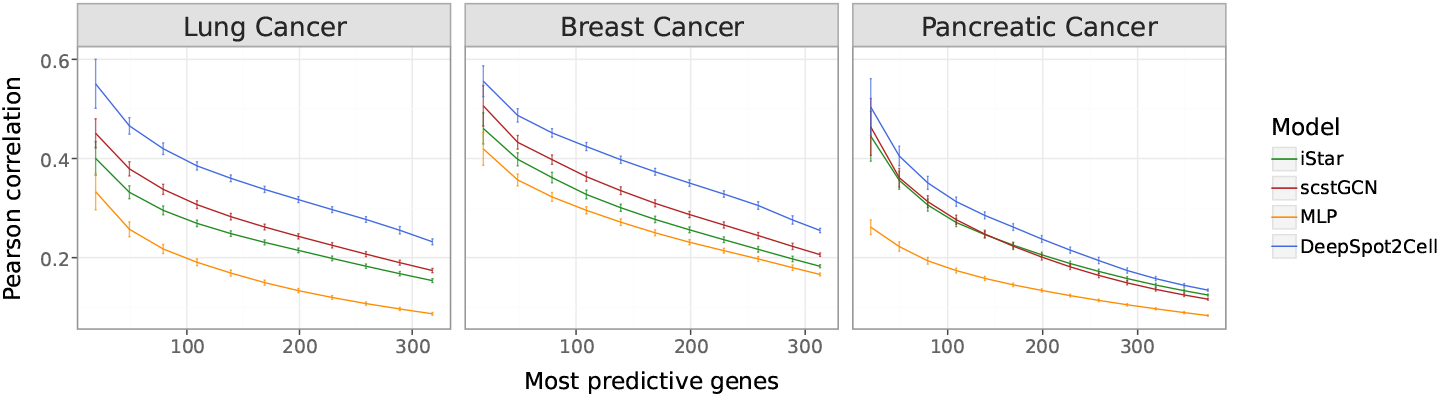
Deconvolution benchmark across lung, breast, and pancreatic cancer datasets. Sets of X most predictive genes for each model on the x-axis are sorted by the descending Pearson correlation on the y-axis. Correlations computed per-gene on the cells from within spots (*IS*_*in*_).

Figure 2 qualitatively illustrates the deconvolution of *MSLN*, a known non-small cell lung cancer marker gene, on slide NCBI867 from the lung cancer dataset. Predictions from iStar (*r* = 0.21) and scstGCN (*r* = 0.30) are noisy and spatially diffuse, whereas DeepSpot2Cell (*r* = 0.45) produces coherent, spatially structured patterns that align better with the ground truth.

IS_*in*_: gene correlation among cells within spots. IS_*out*_: gene correlation among cells outside spots. OOS: gene correlation among cells on hold-out patients, same cohort. OOD: gene correlations on cells from different cohort.

### 4.2 DeepSpot2Cell predicts virtual single-cell spatial transcriptomics from H&E images

Furthermore, the out-of-sample (OOS) and out-of-distribution (OOD) scenarios assess a model’s ability to infer virtual single-cell gene expression for samples unseen during training, simulating its application to novel data from the same or a different cohort, respectively (Table 1).

For example, in the lung cancer OOS scenario, DeepSpot2Cell consistently increased the Pearson correlation by 46%, from 0.24 (best competitor, scstGCN) to 0.35 (DeepSpot2Cell). In the more challenging breast cancer OOD scenario, DeepSpot2Cell improved the gene correlations by more than 50%, demonstrating that it has learned robust single-cell transcriptomic mappings that generalize to unseen histopathological images from other cohorts.

### 4.3 DeepSpot2Cell ablation experiments

Next, we evaluated how specific modeling choices in DeepSpot2Cell contribute to its accuracy in single-cell prediction (Figure 3). We make a few important observations:

1. Leveraging spatial multi-level tissue context through both spot representations and their neighbors improves DeepSpot2Cell’s gene correlations compared with using only the spot and cell or only the cell itself, consistent with previous findings [27].
2. The choice of PFM is important, with Phikon-v2 outperforming UNI and H-Optimus-0, potentially due to Phikon-v2’s multi-resolution training design.
3. A more advanced GRU network for learning the set convolution operation performs worse than simple summation, likely due to cell order sensitivity.

## 5 Discussion & Conclusion

In this work, we propose DeepSpot2Cell, a novel DeepSet neural network that leverages PFM and spatial multi-level tissue context to accurately infer virtual single-cell ST from routine histology images using spot-level supervision. The method’s key innovation is modeling spots as bags of cells: DeepSpot2Cell learns how individual cells contribute to spot-level gene expression and uses these mappings to predict single-cell expression. Further, we curated a newly dedicated benchmark designed for single-cell ST deconvolution and prediction, enabling systematic comparison of models.

Our results demonstrate that DeepSpot2Cell outperforms previous super-resolution models in single-cell deconvolution and prediction across multiple cancer types, even in out-of-distribution settings. The variable performance of other super-resolution models across different scenarios indicates over-fitting to the specific images and transcriptomic spots. In contrast, DeepSpot2Cell gene correlations were consistent, indicating that it has learned general single-cell mappings that could be transferred to unseen images. Notably, both iStar and scstGCN underperformed on OOD breast cancer compared to our MLP baseline, highlighting their limited ability to generalize beyond the training distribution.

In summary, DeepSpot2Cell uses the abundance of spot-based ST data both to augment existing ST cohorts with cell-level resolution and to learn generalizable single-cell transcriptomic mappings, enabling the prediction of single-cell expression profiles from H&E images.

## 6 Limitations & Future work

Several limitations merit acknowledgment. First, the experiments used pseudo-Visium spots derived from Xenium data, which may not fully reflect real Visium measurements. We also relied on available old Xenium datasets with ∼ 300 genes rather than the newer 5k panels. Second, accuracy depends on the quality of cell segmentation, as errors in segmentation propagate to expression assignments. In our benchmark, we relied on Xenium ground-truth nuclei locations, but in practice, these must be inferred computationally, requiring accurate nucleus detection. Improving and integrating these methods is essential for single-cell resolution in real-world settings. Finally, this work motivates further research on the utility of the virtual cells in downstream biological applications, including identifying genes that correlate with tissue architecture and those that cannot be reliably predicted.

## A DeepSpot2Cell training details

DeepSpot2Cell was trained on an NVIDIA RTX 4090 using the Adam optimizer with a learning rate of 10^−4^ and a batch size of 256 spots. Early stopping was used based on validation loss. Dropout with rate 0.3 was applied to the *ϕ* and *ρ*_gene_ MLPs to reduce overfitting. We optimize the model by minimizing the mean squared error (MSE) loss between predicted and observed spot expressions:

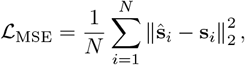

where *N* is the number of spots. The code is available at https://github.com/ratschlab/DeepSpot2Cell. Computational data analysis was performed at Leonhard Med (https://sis.id.ethz.ch/services/sensitiveresearchdata/), a secure trusted research environment at ETH Zurich.

## B Pathology foundation models

Tile embeddings were extracted from pretrained pathology foundation models (PFM), with their weights obtained from Hugging Face.

1. UNI: https://huggingface.co/MahmoodLab/UNI
2. Phikon v2: https://huggingface.co/owkin/phikon-v2
3. H-optimus-0: https://huggingface.co/bioptimus/H-optimus-0

We benchmarked their performance to assess their contribution and found that Phikon v2 produced more accurate expression predictions relative to the ground truth (Figure 3). To isolate the effect of the PFM in our benchmarks, we kept Phikon v2 fixed as the underlying pathology model in DeepSpot2Cell and scstGCN, ensuring fair comparability. Notably, these models are PFM-agnostic in practice, allowing the pathology foundation model to be replaced with a more suitable one depending on tissue characteristics.

## C MLP baseline

The MLP baseline is a two-layer network designed to isolate the contributions of DeepSpot2Cell’s core components: (1) spatial multi-level context integration, and (2) the DeepSets architecture, which handles variable numbers of cells per spot. This baseline provides a direct strategy for inferring single-cell expression from H&E images using PFM features, serving as a reference for evaluating the trade-off between architectural complexity and predictive performance.

Training followed the procedure described in Appendix A, with the MLP optimized to predict spot-level expression from spot-tile PFM features. During inference, cell-level PFM features were provided, and the outputs were interpreted as cell-level predictions.

## D Super-resolution models

Super-resolution methods were trained using default hyperparameters from their respective official implementations.

### D.1 Code availability

1. scstGCN: https://github.com/wenwenmin/scstGCN
2. iStar: https://github.com/daviddaiweizhang/istar

### D.2 Superpixel map expression post-processing details

These methods generate continuous high-dimensional expression grids corresponding to the original patch regions: iStar produces 16×16 node grids while scstGCN outputs 14×14 grids. During evaluation, to obtain cell-level predictions from the grid outputs, cell bounding boxes were manually downscaled to grid coordinates and intersected with the grid nodes. Then, cell expression values were computed as the average of all nodes intersecting with each cell’s downscaled bounding box. The resulting cell-level predictions were then normalized to 10,000 counts per spot, followed by log1p transformation.

## E Data

We utilized organ-specific Xenium datasets, focusing on three cancer types for which sufficient high-quality samples were available: 20 lung [28], 7 breast [29, 30], and 2 pancreatic [31, 32] cancer samples (Table 2). We downloaded the datasets using the HEST-1k preprocessing pipeline [20].

## F Data preprocessing

### F.1 Pseudospots definition

10x Visium spots contain a tissue area of 55µm, capturing between 1-10 cells, depending on the cell size [4]. To account for the cell size variability [25], cells were considered inside the spot if their nucleus fell at least 10µm within the spot boundary, otherwise considered outside the spot. While this approach may be suboptimal—since cancer cells are generally larger [34] and their membranes could extend beyond the spot boundary—we calculated with this setup that the average number of cells per spot across the three datasets to be between 1-10 cells (Figure 4), which alligns with 10x Visium reported characteristics [4].

Specifically, H&E images were available at 20x magnification and were divided into non-overlapping 224×224 pixel tiles. Each tile contained a central circular pseudospot with a 160-pixel diameter, and spot centroids were spaced 224 pixels apart. Spot-level expression was computed as the sum of all cells located within the pseudospot. The distribution of cell counts per spot across all datasets is shown in Figure 4.

### F.2 Gene expression preprocessing and quality control

Gene counts preprocessing followed a standardized pipeline. Genes expressed in fewer than 20 cells across the sample, as well as blank and negative control genes, were removed. For normalization, spot-level counts were scaled to sum to 10,000 transcripts per spot and subsequently log1p-transformed. Cell-level normalization was performed based on spatial context (inside or outside a spot): cells within a spot were normalized to sum to 10,000 counts, whereas outside-spot cells were normalized using the total counts of the inside-spot cells from the corresponding spot. This strategy ensured that outside-spot cells did not affect the normalization of inside-spot cells, while still undergoing consistent preprocessing.

### F.3 Feature extraction

Individual cell tiles were defined as the smallest square that fully contains the segmented area of the cell, which was obtained by CellViT [26] and was provided with the HEST-1k dataset. Cell and spot tiles were transformed in accordance with the recommended preprocessing of each particular pathology foundation model.

## G Evaluation details

Model performance was assessed using per-gene Pearson correlation.

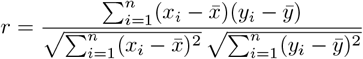

where *x*_*i*_ denotes the predicted value for gene *i, y*_*i*_ denotes the observed value for gene *i*, 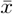and 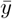 are the mean predicted and observed values, respectively, and *n* is the number of cells.

The Pearson correlation measures how well the predicted values agree with the observed values across samples for each gene. Cross-validation employed patient-level data partitioning to ensure validation splits contained only samples from distinct patients. The number of folds was set to *min(5, n_patients)*. We bootstrapped 10,000 times from the median Pearson correlation across the test folds and reported the resulting median Pearson correlation along with its standard error.

## H Ablation details

All ablations were compared on the lung cancer dataset using the area under the Pearson correlation gene curve computed on cells from within spots (IS_*in*_), as shown in Figure 3.

## I Extended evaluation results

